# Emergent smartphone temporal structures reflect cognitive constraints

**DOI:** 10.64898/2026.04.05.716589

**Authors:** Enea Ceolini, Guido Band, Arko Ghosh

**Affiliations:** QuantActions AG, Zurich Switzerland; Institute of Psychology, Leiden University, Leiden, The Netherlands

## Abstract

Fine-grained temporal structures emerge in smartphone behavioral recordings over multi-day periods. Complex systems research suggests that emergent temporal structures reflect underlying resource constraints of the system. Here we test whether cognitive abilities measured through speeded tasks (spanning fractions of a second) are reflected in emergent smartphone temporal structures spanning days, revealing how cognitive resource limitations shape naturalistic behavior. We analyzed smartphone tap interval patterns accumulated over several days and used decision tree regression models to predict performance in simple and choice reaction time tasks from these patterns. Simple reaction time was poorly predicted (R^2^ = 0.003), indicating that basic sensorimotor constraints play only a marginal role in shaping real-world behavioral timing. In contrast, choice reaction time was moderately predictable (R^2^ = 0.4), demonstrating that higher-order cognitive constraints prominently influence naturalistic temporal organization. Notably, while task performance operates at sub-second timescales, predictive temporal patterns in smartphone behavior spanned milliseconds to several seconds and was accumulated over days, revealing the broad, multi-scale influence of cognitive resource constraints on everyday behavior. Both predicted and measured choice reaction times showed age-related decline, but the decline was more pronounced in predicted values, suggesting that age-related cognitive changes may be amplified in naturalistic contexts. These findings demonstrate that emergent temporal structures in smartphone use can reveal how cognitive processes measured using speeded tasks manifest, or fail to manifest, in real-world behavior. Complex-systems approaches can bridge laboratory and naturalistic assessments of cognition, revealing which cognitive processes meaningfully constrain real-world behavior.

## Introduction

Human behavior in naturalistic settings appears complex and variable, and behavioral outputs are apparently unpredictable. However, the timing of human activities, from mail correspondence to telephone calls to website visits, exhibits emergent temporal patterns such as scale-invariant bursts, suggesting underlying organizational principles (Barabási, 2005; Clauset et al., 2009; Malmgren et al., 2008; Vázquez et al., 2006). A hypothesis from complex systems research proposes that such patterns reflect resource constraints that shape the system’s output, analogous to how server response patterns reveal bandwidth limitations (Adler et al., 1998; Li et al., 2014; Vázquez et al., 2006). Supporting this framework, smartphone touchscreen tap intervals exhibit fractal-like temporal organization spanning ∼1 second to several hours, with bursts of rapid interactions interspersed with longer pauses (Pfister and Ghosh, 2020). Critically, the scaling properties of this fractal-like organization correlates with individuals’ fastest tapping speeds in the sub-second range, suggesting that basic timing constraints propagate across temporal scales to shape multi-hour behavioral dynamics. This constraint-based framework gains convergent support from psychology (Salthouse, 1996): performance on speeded tasks, which probe fundamental cognitive and sensorimotor capacity operating at millisecond timescales, correlates with diverse real-world outcomes including longevity, cognitive decline, and the conduct of instrumental activities of daily living (Bielak et al., 2010; Deary and Der, 2005a; Jakobsen et al., 2011; Steinborn et al., 2016). However, these (mostly weak) associations are established through self-reported functional assessments or distal health outcomes rather than direct quantification of naturalistic behavior, leaving the actual manifestation of the cognitive constraints in everyday behavioral patterns unexplored (Sbordone, 2001).

A novel perspective on smartphone behavior enables us to address how cognitive constraints manifest in emergent behavioral patterns (Duckrow et al., 2021). Essentially, we assume that real-world behavior unfolds as a continuous flow of neurobehavioral states whose complex trajectories cannot be directly observed or interpreted. Inspired by Poincaré sections used to study complex dynamical systems (Holmes, 1990), we treat the smartphone touchscreen as a Poincaré section through this continuous neurobehavioral flow. Each touch on the screen represents a moment when the trajectory intersects this section, and the intervals between consecutive touches quantify the dynamics of this flow (Fig. 1). We characterize these intervals using joint-interval distributions (JIDs)(Brennan et al., 2001; Duckrow et al., 2021; Rodieck et al., 1962): by plotting each inter-touch interval against its successor, analogous to return maps in dynamical systems (Holmes, 1990), we construct a two-dimensional behavioral landscape from accumulated observations. This behavioral map can be used to reveal how cognitive resource limitations that constrain neurobehavioral state dynamics manifest in the temporal organization of real-world behavior.

**Figure 1.**
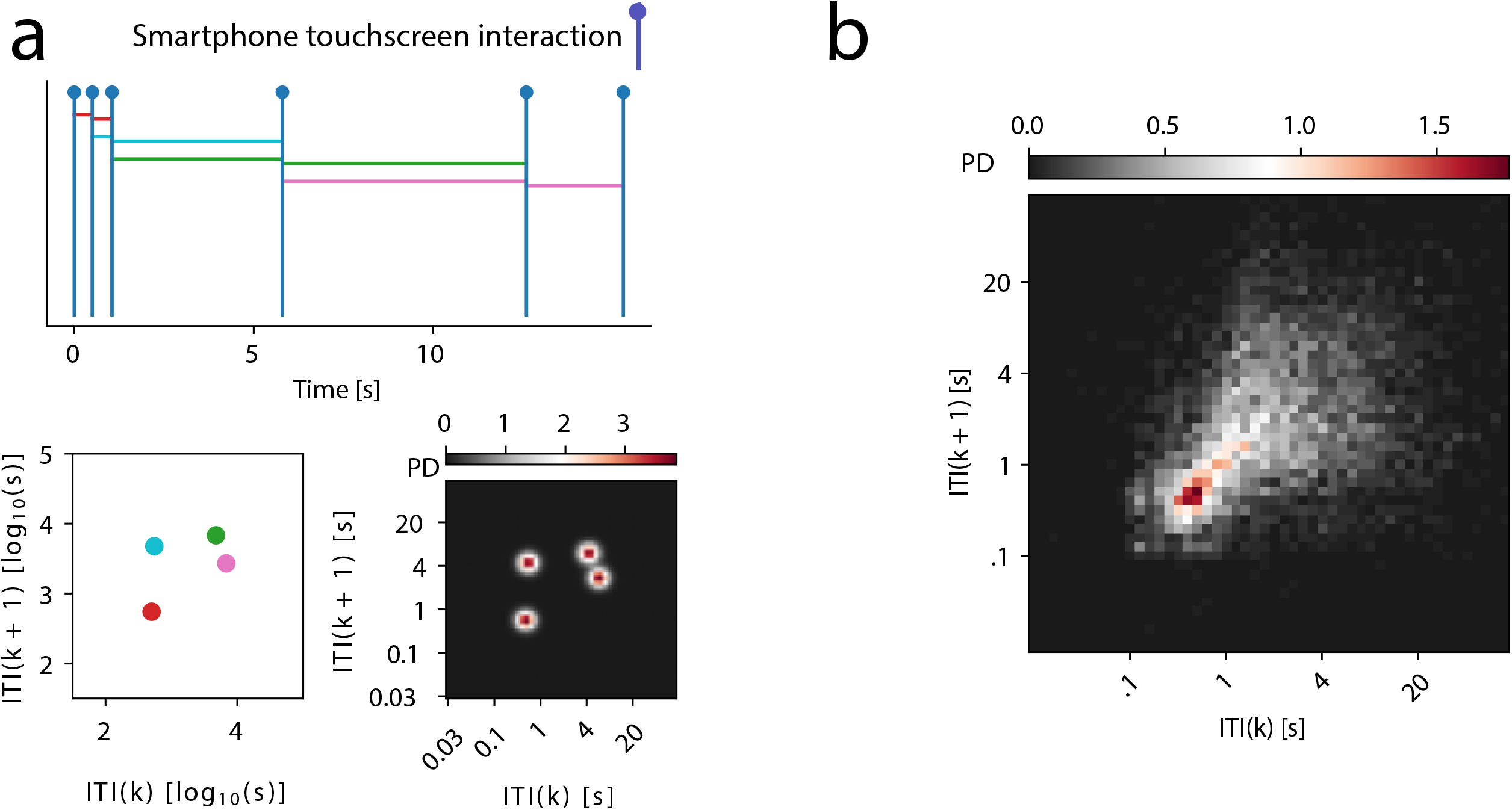
Smartphone behavioral maps based on next-interval dynamics of touchscreen interactions. (a) Schematic illustration of joint-interval distribution (JID) construction. A time series of touchscreen interactions (top) generates inter-touch intervals (ITIs). The JID is constructed by plotting each interval *k* against its immediate successor *k+1*, creating a two-dimensional return map of behavioral dynamics. (b) A JID from an individual participant where the data was accumilated over a period of 4 days. PD, probability density. ITI, inter touch interval.

These low-dimensional smartphone behavioral maps are sensitive to multiple factors operating across timescales. When constructed hourly, they oscillate systematically with circadian and weekly rhythms, as well as other multi-day oscillations (Ceolini and Ghosh, 2023). These hourly maps also reveal signatures of epileptic neural activity (Duckrow et al., 2021). When accumulated over a day or longer, they reflect chronological age in healthy individuals, isolated clinical events such as seizures, and accelerated aging in disease states (Ceolini et al., 2024, 2022a; van Nieuw Amerongen et al., 2024). Mass-univariate analyses have extended this framework by correlating regions of the behavioral map with cognitive task performance (Ceolini et al., 2022b). Critically, choice reaction time; engaging executive control and sensorimotor processes, correlated with large distributed map regions, while simple reaction time; primarily sensorimotor, correlated with focal clusters of rapid touches (Ceolini et al., 2022b). However, this correlational approach treats map regions independently, potentially missing complex interdependencies and not simply linear relationships, and does not demonstrate predictive generalization to new individuals.

Machine learning provides a means to test whether cognitive processes measured by laboratory tasks can be predicted from the emergent smartphone behavioral maps. We focused on two widely used indices: simple reaction time (SRT), which primarily captures sensorimotor processing speed, and choice reaction time (CRT) which engages executive functions including response selection and cognitive control. While CRT shows moderate correlations with age and general intelligence, SRT shows weaker associations with the same variables (Deary et al., 2011; Deary and Der, 2005b). Both demonstrate clinical sensitivity in neurological diseases (Eckner et al., 2014; Pirozzolo et al., 1981). Here we trained decision tree regression models to predict individual reaction times from smartphone JID features accumulated over multiple days. We leveraged interpretability tools to isolate which JID features drove predictions, revealing that predictive patterns spanned all time scales of the JID (from milliseconds to ∼30 s). We found that CRT was moderately predictable while SRT showed negligible predictive power, demonstrating that higher-order cognitive constraints, but not basic sensorimotor speed, systematically shape naturalistic behavioral timing.

## Results

### Smartphone touchscreen dynamics predicts reaction time

Participants completed simple and choice reaction time (CRT) tasks during laboratory testing sessions. We used leave-one-participant-out cross-validation to predict individual reaction times from smartphone JID features and chronological age (Fig. 2). We tested multiple temporal windows and found that behavioral data from the 4 days immediately preceding each testing session yielded optimal predictions based on a composite performance metric (F-score; see Methods). This yielded 360 models (one per participant), which we evaluated at two levels: participant-level predictions using median performance across repeated sessions (n=360), and session-level predictions including all testing occasions (n=2,633).

**Figure 2.**
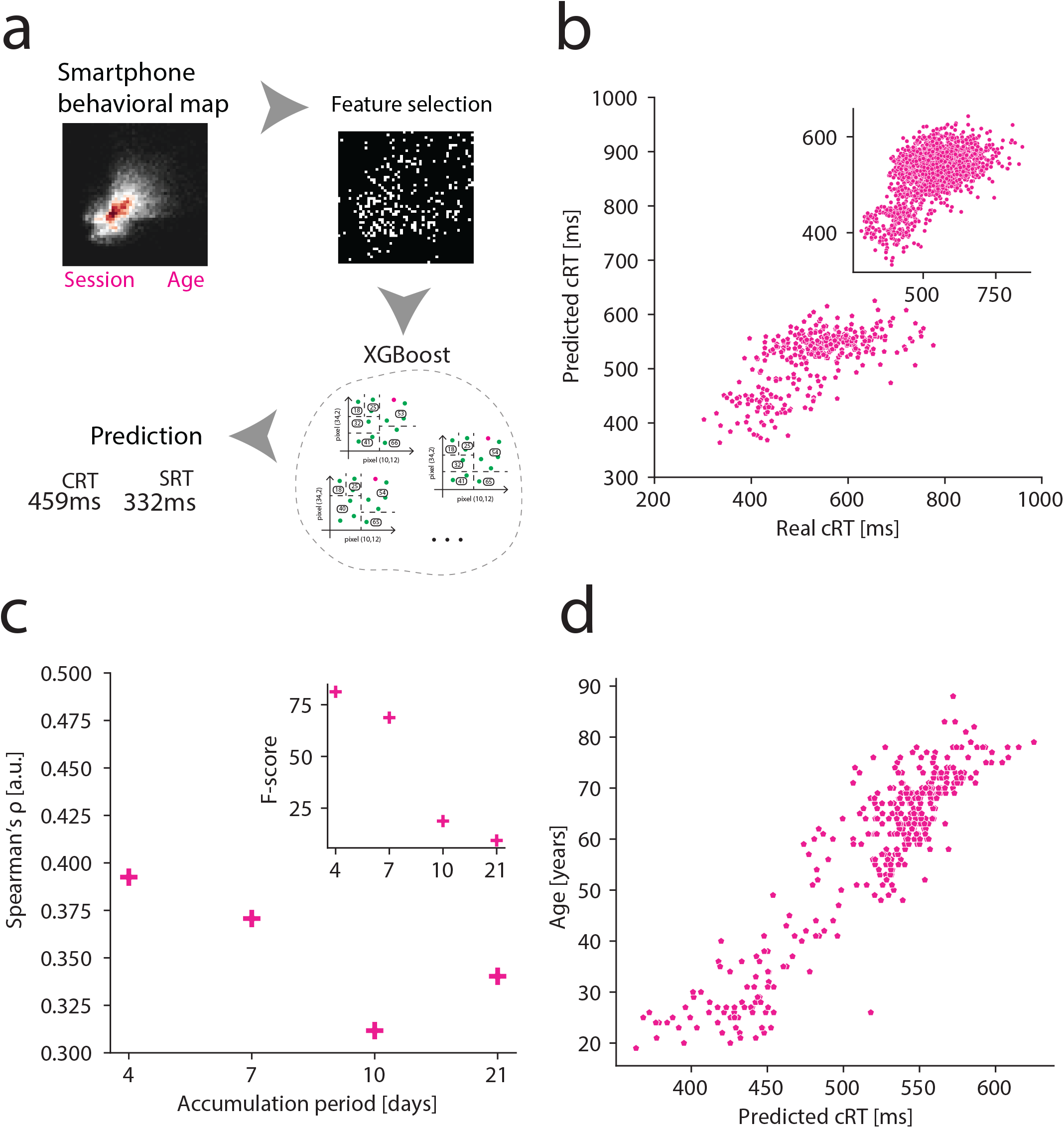
Smartphone behavioral maps predict choice reaction time. (a) Analysis pipeline. Joint interval distributions (JIDs) from multi-day smartphone use were used to predict reaction time performance via XGBoost regression with feature selection. (b) Predicted versus measured choice RT at session level (top) and participant level (bottom). (c) Model performance by accumulation period (Spearman’s ρ). The 4-day window preceding testing yielded optimal predictions (F-score combining Spearman’s ρ, R^2^, MAE, RMSE, inset). (d) Age-related decline in predicted choice RT from models excluding chronological age.

At the participant-level, CRT was substantially predictable from smartphone behavioral patterns (t(359) = 15.83, p = 3.8 × 10^−43^, R^2^ = 0.41, simple linear regression), while SRT showed essentially no relationship (t(359) = 1.05, p = 0.295, R^2^ = 0.003). This striking dissociation persisted at the session-level: CRT remained significantly predictable (t(2632) = 30.66, p = 1.0 × 10^−176^, R^2^ = 0.26), while SRT, despite achieving statistical significance due to large sample size, showed negligible explained variance (t(2632) = 9.80, p = 2.8 × 10^−22^, R^2^ = 0.04). These findings demonstrate that naturalistic smartphone behavioral dynamics reflect higher-order cognitive constraints measured by CRT, but not basic sensorimotor processing speed measured by SRT. Next, we address how these predictions alter with age.

### Predicted reaction time declines with age

Our sample spanned the adult lifespan (18 to 86 years of age). Given that reaction times slow with age, we examined whether this age-related decline was evident in smartphone-predicted RT values. Critically, we used multivariate regression to test whether predicted RT values provide distinct information about age beyond what is captured by laboratory-measured RT values. Towards this, we raised new sets of leave-one-out models, but the chronological age was withheld (n = 363). We then used a multi-variate regression model with the real and predicted RT (median across sessions) as independent variables and chronological age as the dependent variable (Fig. 2). For the choice reaction time, the full model was strongly related to age (*F*(2, 360), *p* = 7.10 x 10^-73^, *R*^2^ = 0.603, multi-variate regression). Both the predicted and real CRT independently contributed to this relationship: *β*_predicted_ = 0.17 (SE = 0.014) *t* = 12.47, *p* = 6.5 × 10^−30^, *β*_real_ =0.07 (SE = 0.01) t = 9.75, *p* = 4.5 × 10^−20^. Notably, the coefficient for predicted RT was more than twice as large as for measured RT, indicating that smartphone-derived behavioral patterns capture age-related cognitive changes substantially more strongly while also providing complementary information.

### Behavioral features underlying CRT predictions

To identify which smartphone behavioral patterns predicted CRT, we applied SHAP-based feature importance analysis to each leave-one-out model (see Methods). For each bin in the two-dimensional JID map, we quantified how frequently it was selected across all 360 models (Fig. 3). JID features representing sub-second tap intervals were consistently selected and clustered in specific regions of the behavioral map, while features representing longer intervals (>1 second) showed spatially distributed selection across the map without clear clustering. This pattern indicates that CRT, which operates at sub-second timescales, relates to specific, consistent patterns of rapid smartphone interactions, whereas its relationship to slower behaviors is more diffuse.

**Figure 3.**
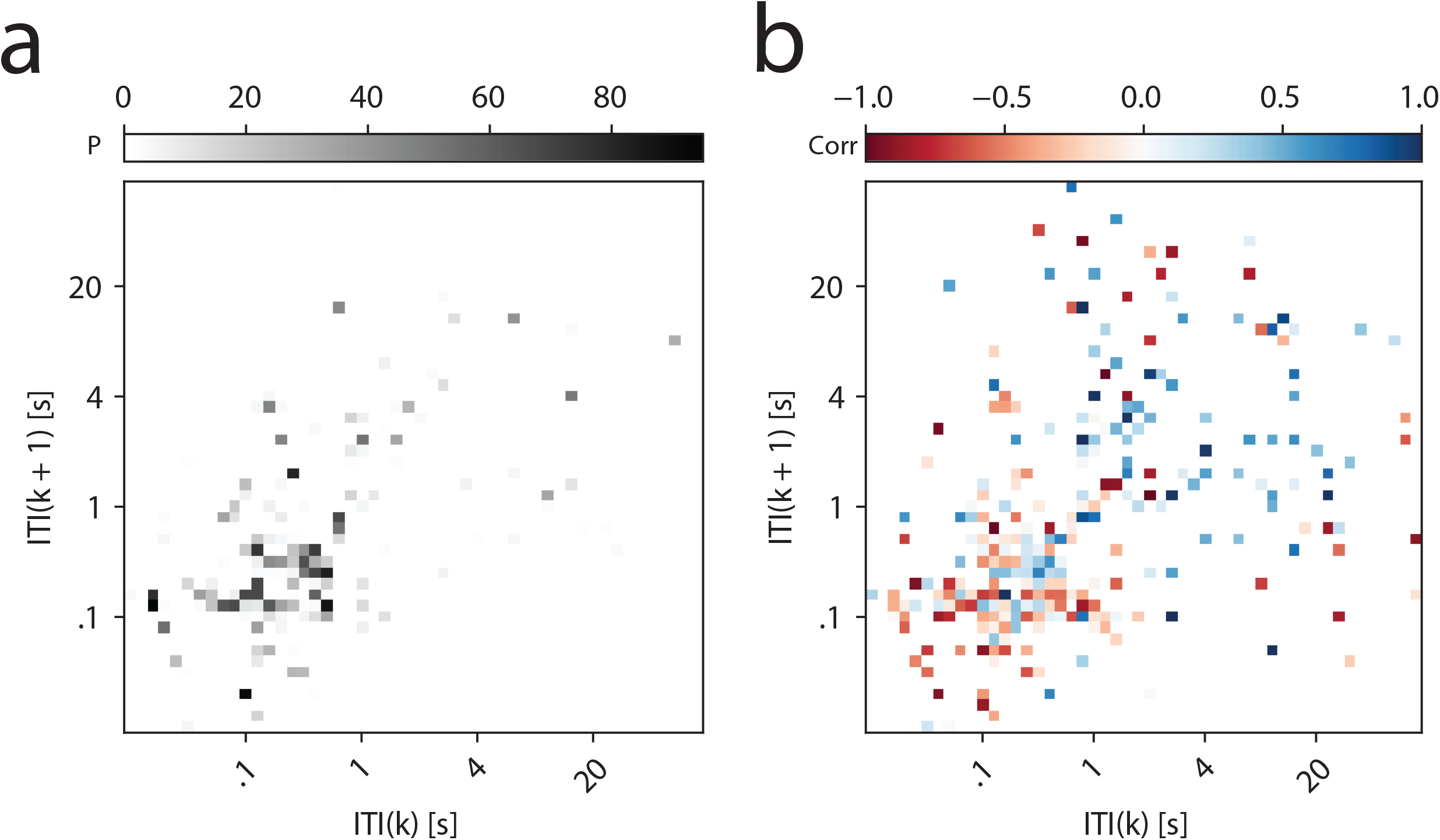
Behavioral features underlying the prediction of choice reaction time (CRT). (a) Frequency of feature selection across all models trained using a leave-one-participant-out cross-validation strategy. Darker pixels indicate features that were consistently selected across participants, reflecting their importance in predicting CRT (b) Directional contributions of individual features expressed as population level correlation coefficients (based on all the leave one participant out models) at a given two dimensional bin (feature) between behavioral probability and SHAP values. For instance +1 indicates strong directly proportional link: more the behaviour slower the reaction time, and -1 indicates an inversely proportional link: more the behaviour faster the reaction time.

We next estimated the directional impact of selected features, whether they predicted slower or faster CRT based on SHAP values (Fig. 3). Lower frequencies in clustered sub-second intervals and higher frequencies in spatially distributed features (spanning both sub-second and longer intervals) both predicted slower CRT. Conversely, higher frequencies in clustered sub-second intervals and lower frequencies in distributed features predicted faster CRT. However, the relationship was not simply divided between sub-second versus supra-second behaviors: some sub-second intervals predicted slower CRT while others predicted faster CRT, depending on their location in the behavioral map. Despite this complexity, a discernible core pattern emerged wherein very rapid consecutive interactions (<0.3s intervals) most consistently predicted faster cognitive performance.

## Discussion

Our findings distinguish the manifestation of two widely used speeded cognitive tasks, simple reaction time (SRT) and choice reaction time (CRT), in emergent behavioral patterns. It is well established that SRT and CRT represent different cognitive processes (Donders, 1969; Suarez et al., 2021), best reflected in the way decision-making is modeled in computational approaches of RT distributions (Ratcliff and McKoon, 2008; Smith, 1995). While previous studies showed dissociations of SRT versus CRT at the group level and in laboratory settings (Godefroy et al., 2010) our work demonstrates that this distinction extends to their manifestation in real-world behavior. Emergent temporal structures in smartphone use accumulated over multiple days strongly reflect CRT performance but show negligible relationship to SRT. This dissociation reveals two key insights: First, the cognitive processes underlying CRT, including response selection and cognitive control, pervasively influence the temporal organization of everyday behavior. Second, basic sensorimotor speed measured by SRT plays minimal role in shaping naturalistic behavioral timing, highlighting its limited relevance for understanding real-world cognitive function.

Our analysis traced cognitive processes through emergent temporal structures in naturalistic behavior. Three mechanistic interpretations merit consideration. First, cognitive constraints may directly shape behavioral organization: CRT-related processes systematically structure smartphone timing, analogous to how bandwidth constraints shape server outputs in complex systems (Pfister and Ghosh, 2020; Vazquez, 2007). Simple sensorimotor speed, in contrast, shows minimal influence. Second, use-dependent plasticity could operate in reverse: smartphone behavior might shape CRT processes. While smartphone use has been associated with neural plasticity, existing evidence focuses on sensorimotor rather than executive domains (Balerna and Ghosh, 2018; Ghosh, 2021; Gindrat et al., 2015). Third, common factors, including stable traits (age, personality) or fluctuating states (arousal, mood), could independently influence both RT performance and behavioral patterns. The recency effect, wherein the 4 days immediately preceding testing most strongly predict RT, provides crucial constraints. This temporal specificity rules out an exclusive role of stable trait factors and indicates that any operative mechanism must function on timescales of days or less. Whether this reflects cognitive constraints dynamically shaping behavior, rapid plasticity, or short-term state fluctuations remains to be fully determined.

The behavioral features predicting CRT spanned both sub-second and supra-second timescales. This is consistent with evidence that cognitive processes underlying speeded decisions also support slower choices (Krajbich et al., 2015), suggesting that executive control and response selection operate across temporal scales. This multi-scale manifestation may explain why age-related decline was more pronounced in predicted CRT compared to measured CRT values. Smartphone behavior samples CRT-related cognitive processes across diverse naturalistic contexts—from rapid typing to deliberate browsing—providing multiple independent observations of the same underlying constraints. This broader sampling may amplify age-related signals that are less apparent in brief laboratory assessments. These findings align with established patterns showing that CRT, but not SRT, demonstrates strong age-related decline and disease sensitivity (Bielak et al., 2010; Deary et al., 2011; Myerson et al., 1990; Pirozzolo et al., 1981), likely reflecting the more widespread engagement of executive control processes across behavioral contexts.

Our findings demonstrate that emergent smartphone behavioral structures reflect specific cognitive constraints: CRT-related executive processes and other relatively slow cognitive processess pervasively shape naturalistic behavior, while basic sensorimotor speed does not. This bridges complex systems approaches to behavior with individual differences in cognitive function, revealing how latent processess underlying laboratory-based measures manifest,and potentially amplify, in real-world contexts. Practically, this enables passive, continuous cognitive monitoring through naturalistic smartphone use, overcoming the logistical constraints of traditional laboratory testing. This approach may prove particularly valuable for early detection and longitudinal monitoring of cognitive decline, offering ecologically valid assessment that complements conventional neuropsychological methods.

## Methods

### Participants

We recruited healthy participants (self-reported; health status assessed using SF-36) through our online laboratory platform, with recruitment aided by the Dutch Brain Registry (Zwan et al., 2021). Portions of this dataset have been reported previously (Ceolini et al., 2024, 2022b; Ceolini and Ghosh, 2023). We recruited 720 volunteers, of whom 642 installed the smartphone data collection app (see below) and provided demographic information (235 males, 396 females, 11 prefer not to say; ages 16–86 years). The present analysis includes 360 participants who completed at least one reaction time testing session and accumulated at least 100 touchscreen interactions during the corresponding 4-day window. All procedures were approved by the local ethics committee at Leiden University.

### Smartphone data collection app

Participants installed the TapCounter app (QuantActions AG, Zurich, Switzerland) from the Google Play Store on their personal Android smartphones. The app passively recorded all touchscreen interactions with millisecond-precision timestamps and associated app identifiers (Balerna and Ghosh, 2018). Data were automatically uploaded to a secure cloud platform without interfering with normal device use. For the present analysis, we used only tap timestamps to construct joint-interval distributions; app identity information was not analyzed.

### Cognitive tests

Participants completed the Deary-Liewald simple and 4-choice reaction time tasks (Deary et al., 2011) via PsyToolkit (Stoet, 2017), as described previously (Ceolini et al., 2022b). Testing was conducted online on participants’ personal computers (keyboard required; participants confirmed appropriate setup before beginning). Tasks were available in English or Dutch. Participants were invited to complete monthly sessions over 53 months, with participation ranging from 1 to over 100 sessions per individual. Each session included 8 shared practice trials followed by 25 SRT trials and 50 4-CRT trials. Inter-trial intervals varied randomly between 1-3 seconds. Only correct trials with RTs ≤1000 ms were analyzed (eliminating a total of 8 trials across the sample). Median RT values for each task and session were used in all analyses.

### The joint interval distribution (JID)

Inter-tap intervals (ITIs) were analyzed using joint-interval distributions (JIDs) to capture sequential timing structure (Duckrow et al., 2021). Only ITIs during screen-on periods were included; screen-off intervals were excluded. Each JID represented the two-dimensional relationship between consecutive ITIs (interval *k* vs. interval *k+1*). ITIs were log_10_-transformed and discretized into 50×50 bins spanning 10^0.5^ to 10^4.5^ ms (∼3 ms to ∼30 seconds), covering the 1st-99th percentile range across participants. Joint probability densities were estimated using scikit-learn (Pedregosa et al., 2011) yielding 2,500 features per participant. We compared JIDs constructed from multiple temporal windows (4, 7, 10, and 21 days) preceding each cognitive test to identify the optimal accumulation period. To identify the optimal temporal accumulation window, we compared JIDs constructed from the 4, 7, 10, and 21 days preceding each test session, yielding 360 (2,633 sessions), 361 (2,648), 361 (2,660), and 363 (2,687) participants respectively. Sample sizes increased slightly with longer windows as more sessions met the minimum data requirement (≥100 taps).

### Decision tree regresion models

Reaction times were predicted using XGBoost gradient-boosted decision trees(Chen and Guestrin, 2016). Predictors included 2,500 JID features (50×50 bins) from the 4-day pre-test window and session number. Observations were weighted by tap count (higher counts indicate more reliable JID estimates). We trained models both with and without chronological age as a predictor to assess behavioral feature contributions beyond age-related variance. We used leave-one-out cross-validation, training 360 models (one per participant) with conservative hyperparameters (learning_rate=0.01, max_depth=4; other parameters at defaults). Model performance was evaluated via linear regression of predicted versus observed values. Feature importance was assessed using SHAP values (Lundberg and Lee, 2017), which quantify each feature’s directional contribution to predictions. For each JID bin, we calculated selection frequency across models and aggregated directional effects.

### Cross-Validation and Feature Selection

We used leave-one-participant-out cross-validation (LOOCV), training n (∼360) models each on n -1 participants and testing on the held-out participant. Within each training set, nested 10-fold cross-validation with early stopping determined optimal boosting rounds. To reduce overfitting given the limited sample size relative to features (2,500 JID bins), we implemented SHAP-based feature selection for each LOOCV model. After training on all features, we calculated SHAP values across nested CV folds and ranked features by importance. We then retrained using progressively smaller feature subsets (500, 250, 125, 60, 15 features), selecting the smallest subset maintaining ≥90% of full-model performance. This consistently yielded 60 features per model, though specific features varied by training set. Models used pseudo-Huber loss and reaction times were z-scored per training set. Performance was evaluated by pooling all test predictions and computing linear regression statistics.

### Selection of the Optimal Accumulation Period

To identify the optimal accumulation period, we compared four window lengths (4, 7, 10, 21 days) using the model variant with age. We evaluated each using four metrics: Spearman’s ρ, R^2^, MAE, and RMSE. For each metric, we ranked the eight conditions and normalized ranks to a 0-100 scale (inverting for MAE and RMSE where lower is better). The composite performance score was the mean of these four normalized values, equally weighting all metrics. The 4-day window consistently achieved the highest composite scores and was selected for all analyses

## Aknowledgements

This project was funded by Velux Stiftung (Grant No.1283, awarded to A.G.) and postdoc Mobility awarded to Enea Ceolini with Arko Ghosh as host (SNSF no. 199692). We also thank all participants for contributing their time and effort. The authors used Claude (Anthropic, Claude Sonnet 4.5, accessed January 2026) to improve the clarity, grammar, and organization of the manuscript text. The AI was used solely for language editing; all scientific content, study design, data collection and analysis, interpretation of results, and conclusions were developed by the authors who take full responsibility for the accuracy and integrity of the manuscript.

